# AmyloDeep: pLM-based ensemble model for predicting amyloid propensity from the amino acid sequence

**DOI:** 10.1101/2025.09.16.676495

**Authors:** Alisa Davtyan, Anahit Khachatryan, Rafayel Petrosyan

**Affiliations:** Institute of Physics, Yerevan State University, 0025 Yerevan, Armenia; Zaven & Sonia Akian College of Science and Engineering, American University of Armenia, 0019 Yerevan, Armenia

## Abstract

Amyloids are predominantly β-sheet-rich, stable protein structures that can maintain their presence in the human body for multiple years. Amyloid protein aggregates contribute to the development of multiple neurodegenerative diseases, such as Alzheimer’s, Parkinson’s, and Huntington’s, and are involved in different vital functions, such as memory formation and immune system function. Here, we used advanced machine learning and deep learning techniques to predict amyloid propensity from the amino acid sequence. First, we aggregated labeled amino acid sequence data from multiple sources, obtaining a roughly balanced dataset of 2366 sequences for binary classification. We leveraged that data to both fine-tune the ESM2 model and to train new models based on protein embeddings from ESM2 and UniRep. The predictions from these models were then unified into a single soft voting ensemble model, yielding highly robust and accurate results. We further made a tool where users can provide the amino acid sequence and get the amyloid formation probabilities of different segments of the input sequence. Users can access the light version of AmyloDeep through the web server at https://amylodeep.com/, and the full model is available as a Python package at https://pypi.org/project/amylodeep/.

**Graphical abstract:** 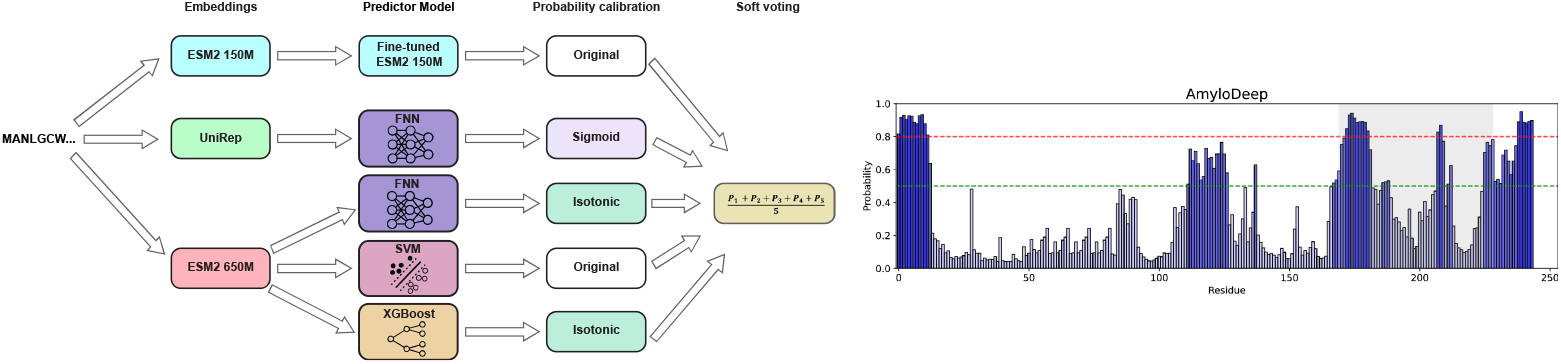

## Introduction

Amyloids are long filamentous structures typically rich with beta sheets (1, 2). Toxic amyloids are involved in many neurodegenerative diseases, while functional amyloids are involved in important physiological functions (3) and can even serve as materials for various applications (4, 5). Despite the recent success of protein structure prediction from its amino acid sequence (6–8), the prediction of the amyloid-forming regions from the amino acid sequences is still an open question. The fact that the same polypeptide chain can fold into a globular native state and, in other cases, can be part of a long amyloid fibril with vastly different structure is one of the demonstrations of the non-uniqueness of sequence to structure correspondence. The identification of the amino acid sequence regions that are prone to amyloid formation is the first important step for their further investigation (9–12). Consequently, the availability of precise and accessible tools for such tasks is of great importance.

Two important developments in recent years contributed to the creation of our tool. First, thanks to the development of new experimental techniques (13, 14) and efficient protocols, the number of experimentally confirmed amino acid sequences that are shown to form amyloids under *in vivo*-like conditions has grown (15–21). It is known that many proteins and peptides can form amyloids under appropriate conditions (4, 22). We, however, are mostly interested in amyloidogenesis under physiologically relevant conditions, which is the primary target of most current methodologies (9, 10, 23–26). Second, the development of novel machine learning (ML) and deep learning (DL) algorithms and the emergence of protein language models (pLMs) have revolutionized the field of protein science (27, 28). pLMs trained exclusively on primary sequence data can learn emergent properties not explicitly provided during training. For instance, these models can infer three-dimensional structural information (28, 29) and enable the *de novo* design of proteins with desired functionalities (30, 31). This is possible since protein sequences are not random amino acid sequences, and models with transformer-based architectures have proven to be able to learn ‘protein language grammar’ (32). Here, we developed a novel, highly accurate pLM-based ensemble model for predicting the amyloid formation probability from the amino acid sequence. We aggregated data from several experimentally verified databases (15–20), used ESM2 (33) and UniRep (34) embeddings, and created five binary classification ML models. These were then combined into a single soft voting ensemble model to provide the final probability values. We then developed a tool that provides the aggregation profile for an amino acid sequence with a user-defined running window size.

## Materials and methods

### Data aggregation

To obtain a unified dataset for amyloid classification, we integrated data from six experimentally verified databases: CPAD 2.0 (https://web.iitm.ac.in/bioinfo2/cpad2/index.html) (15), Waltz-DB 2.0 (http://waltzdb.switchlab.org/) (16), Amyloid Atlas (https://people.mbi.ucla.edu/sawaya/amyloidatlas/) (17), AmyPro (http://amypro.net) (18), AmyLoad (http://comprec-lin.iiar.pwr.edu.pl/amyload/) (19), and AmyloBase (http://bioserver2.sbsc.unifi.it/AmyloBase.html) (20). These resources varied in file formats and annotation detail, requiring parsing and preprocessing pipelines. When available, UniProt identifiers were used as standard keys to combine entries across datasets. We eliminated repeating sequences, removing entries where identical amyloid-forming regions appeared in multiple databases with matching labels, retaining only one copy. We ended up with 2366 sequences, 1237 labeled as amyloid and 1129 as non-amyloid. Data was split into three parts: training set 70 %, validation set 15 % and test set 15 %.

## Models

Fine-tuned ESM2 150M: For the amyloid classification task, we used the ESM2 pLM (33). We used a transfer learning approach wherein the pretrained ESM2 parameters remained frozen while only the classification layer was fine-tuned for binary amyloid/non-amyloid classification. The ESM2 model family includes variants of different sizes, ranging from 8 million to 15 billion parameters. For a comprehensive evaluation, we fine-tuned four variants—ESM2-T6-8M, ESM2-T12-35M, ESM2-T30-150M, and ESM2-T33-650M—using the same training protocol and dataset. All models were initialized with pre-trained weights and fine-tuned with a frozen backbone, updating only the classification head parameters. Among the evaluated variants, the ESM2-T30-150M model consistently demonstrated superior performance for our task, achieving the best trade-off between model capacity and generalization. While the 650M model showed comparable results on training data, it exhibited signs of overfitting and required significantly more computational resources. In contrast, the 150M model provided stable convergence, with higher accuracy and sensitivity. Hence, we selected the 150M model as our final architecture and continued fine-tuning it for additional epochs using early stopping criteria based on validation loss. Hyperparameters such as learning rate, batch size, and dropout rate, as well as the number of epochs, were tuned via grid search within the cross-validation framework. Additionally, we experimented by unfreezing some encoder layers in addition to the classification head of the selected ESM2-T30-150M model. While this setup allowed the model to better capture sequence patterns, it also introduced overfitting. During training, we observed rapid decreases in training loss accompanied by degradation in validation performance, indicating that the model was memorizing the training data rather than generalizing. We tried regularization techniques and hyperparameter tuning, but still didn’t overcome overfitting. Consequently, only the classification layer was fine-tuned, while the core parameters of the pretrained ESM2-T30-150M model remained unchanged.

Unified representation (UniRep) of sequences combined with feedforward neural network (FNN): UniRep is a multiplicative LSTM trained on protein sequences to learn a general-purpose, numerical representation (embedding) of proteins directly from sequence data (34). It generates fixed-length (1900) embeddings for amino acid sequences. We have used the UniRep output embeddings and trained a FNN. It consists of 5 hidden linear layers, with ReLU activation functions, and to prevent overfitting, we incorporated dropout layers with a rate of 0.5. This model is the fastest for training and inference, and is deployed on our web server as a light version.

ESM2 650M combined with three separate ML models: We have used the pre-trained pLM ESM2 650M, to extract the embedding layer outputs (the resulting matrix has a size of a length of amino acid sequence by 1280) and used them as input features for three separate binary classification models. One of these models is a FNN consisting of 5 linear layers with ReLU activations, and to prevent overfitting, we incorporated dropout layers with a rate of 0.2. The second classification model is a support vector machine (SVM) classifier implemented with a linear kernel and regularization parameter C=1. The third classification model is an XGBoost classifier configured with 10 estimators, a maximum tree depth of 3, and a learning rate of 0.1.

Each model was validated using 5-fold cross-validation to ensure robustness and generalization across different data splits. Models’ performances were evaluated using multiple metrics: accuracy, sensitivity, specificity, F1 score, Matthews Correlation Coefficient (MCC), area under the receiver operating characteristic curve (AUC), and average precision (AP), details can be seen in Supplementary Information. Following the experimentation and tuning phase, the final models were trained from scratch on an A100 GPU for about 30 hours.

### Soft voting ensemble model

The predictions of the above-described five models were used to construct an ensemble model with soft voting. We evaluated calibration with reliability diagrams and applied either sigmoidal (Platt) or isotonic calibration, selecting the method per model using the reliability plot, expected calibration error (ECE), accuracy, and specificity (35, 36). As a result, the fine-tuned ESM2 150M and ESM2 650M with SVM were kept as they were, for ESM2 650M with FNN and ESM2 650M with XGBoost, isotonic calibration was applied, and for UniRep with FNN, sigmoidal calibration was applied. The ensemble prediction is the arithmetic mean of the five models’ predicted probabilities.

## Results

### AmyloDeep ensemble model

The schematics of the AmyloDeep ensemble model is presented in Fig. 1. First, the ESM2 and UniRep models process the input amino acid sequence to generate numerical embeddings. These embeddings then serve as input features for a set of downstream predictor models – FNN, SVM, or XGBoost. After the calibration step, the ensemble model prediction probability is calculated as the arithmetic mean of the probabilities from five binary classification models. The ensemble model’s overall performance is better than that of any of its components.

**Figure 1.**
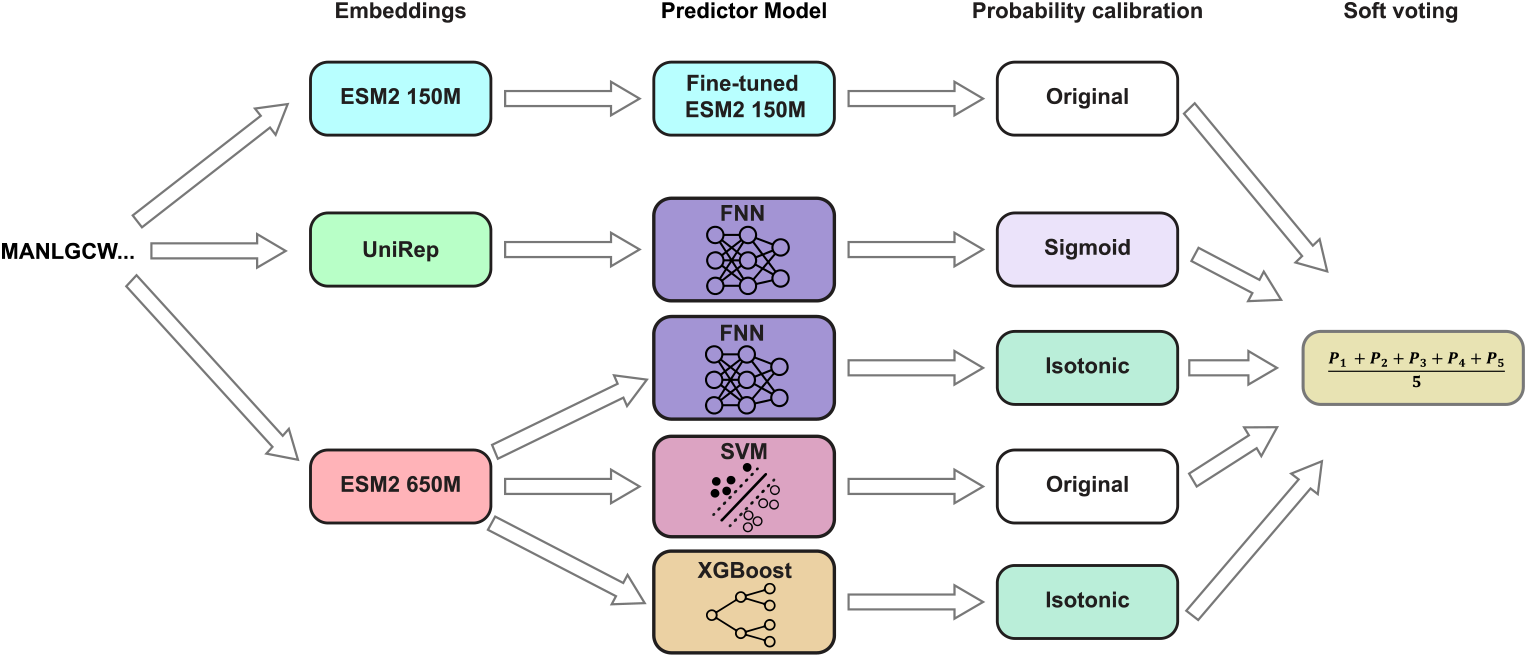
Schematics of the AmyloDeep. The input sequence is sent to ESM2 150M and 650M versions (33) and to UniRep (34), from which the embeddings are sent to the predictor models. Afterwards, the appropriate probability calibration is applied, and the arithmetic mean of the resulting probabilities from five models is calculated.

### Performance

The main performance metrics for the ensemble model are shown in Table 1. Receiver operating characteristic curve (ROC) and Specificity-Sensitivity curve obtained on the test set are presented in Fig. 2A and B, respectively. Overall, this shows a very good performance of our classification ensemble model. However, these are the absolute results, and next, we show the comparisons of AmyloDeep with other similar tools.

**Table 1.**
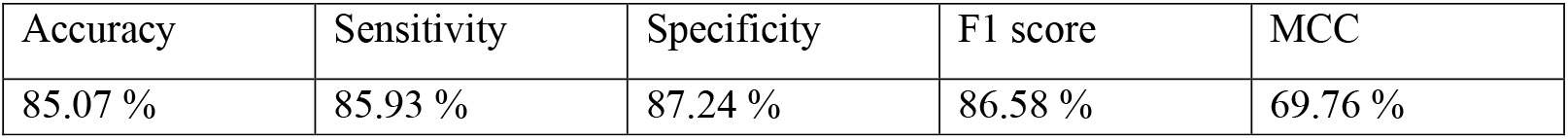
AmyloDeep performance metrics for the amyloid class prediction on the test set.

**Figure 2.**
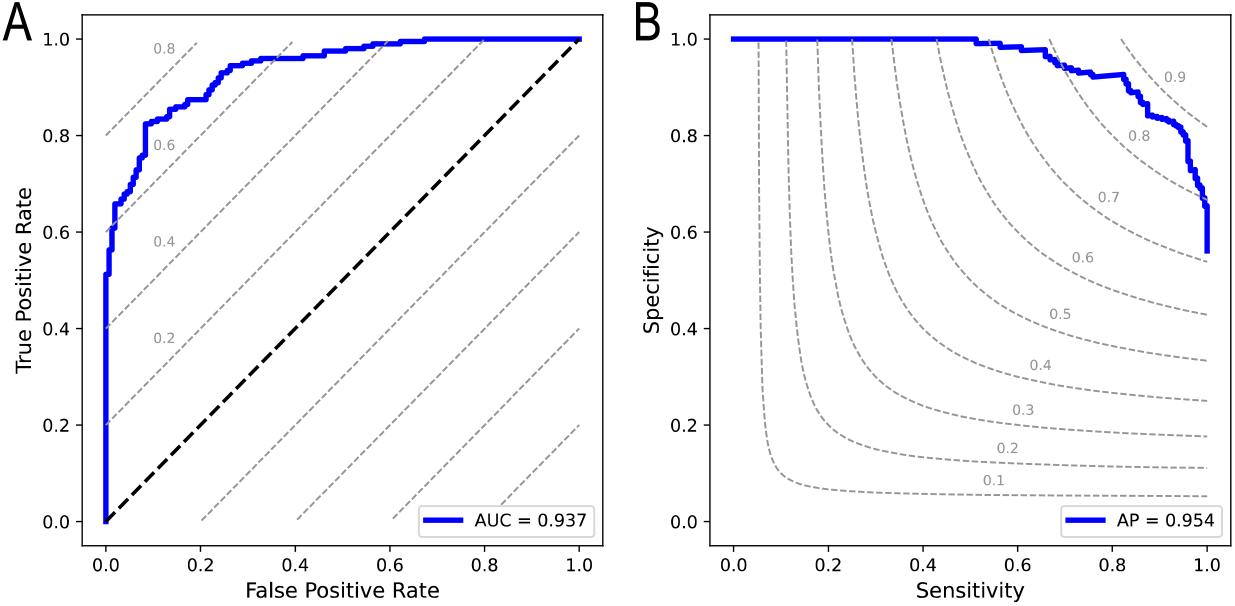
A.ROC curve obtained on the test set with area under the curve being 0.937 TPR·FPR. The dashed lines are iso-performance lines with corresponding TPR – FPR values labeled on them. The black dashed diagonal line corresponds to a random classifier with TPR – FPR = 0. B. Specificity-Sensitivity curve obtained on the test set, average precision is 0.954. The dashed gray lines are iso-F1 score curves with corresponding F1 scores labeled on them.

### Comparison with other methods

In this section, we compare the performance of AmyloDeep with several other methods for amyloid propensity prediction. In Fig. 3, we show the specificity, sensitivity, F1 score, and accuracy of the Cross-Beta predictor (26), AMYPred-FRL (24), and AmyloDeep using sequences from our test set that had a length of 15 amino acids or more, since the Cross-Beta predictor provides predictions only on sequences that are at least 15 amino acids long. AmyloDeep overperforms both models on all of these metrics.

**Figure 3.**
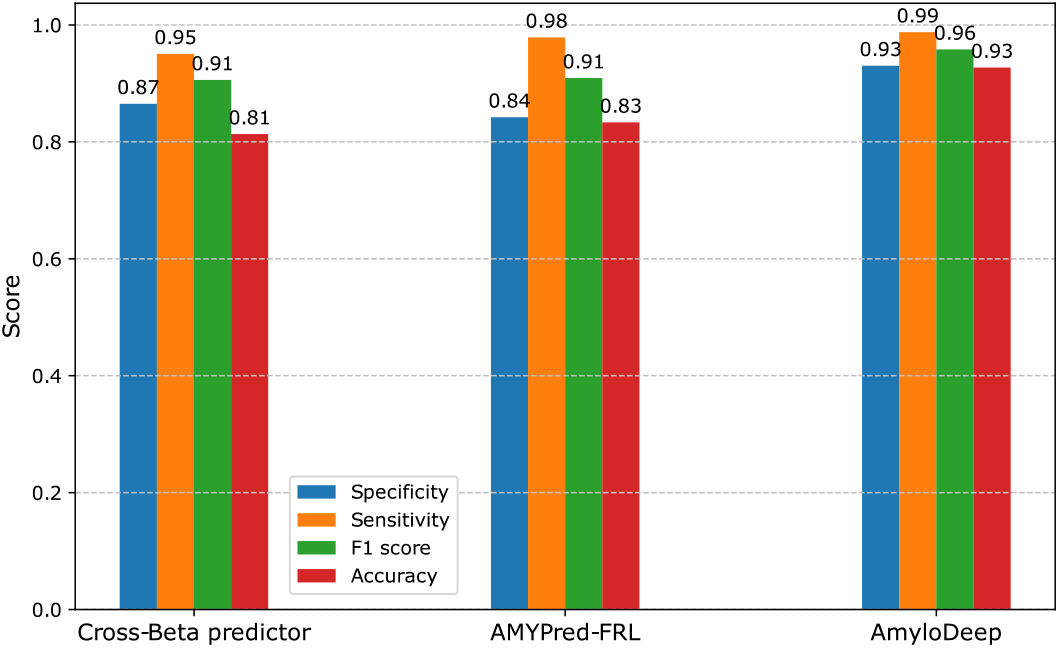
Comparison of AmyloDeep with recently proposed ML-based methods, Cross-Beta predictor (26), and AMYPred-FRL (24). We selected sequences longer than 15 amino acids from our test set since the Cross-Beta predictor provides predictions only on sequences that are at least 15 amino acids long.

Based on our ensemble model, we developed a tool that provides an aggregation profile for a given amino acid sequence. Here, users can provide the amino acid sequence, specify the running window size (default is 10), and obtain an aggregation profile. The light version is available through the web server at https://amylodeep.com/, and the full model is available as a Python package https://pypi.org/project/amylodeep/. Next, in Fig. 4, we compare aggregation profiles predicted by AmyloDeep and two widely used tools: ZipperDB and AGGRESCAN (10). For ZipperDB, if Rosetta energy is below -23 kcal/mol, then it predicts higher than average aggregation propensity, and if Rosetta energy is below -25 kcal/mol, then we have aggregation-prone regions. Because AmyloDeep returns an amyloid-formation probability, we use two decision thresholds: regions with a probability above 0.5 are labeled amyloid-prone, and those with a probability above 0.8 are labeled high-confidence amyloid-prone. In Fig. 4A, we show the aggregation profile for Human PrP (UniProt: P04156) with the gray shaded area corresponding to the amyloid fibril spanning residues 170−229 (PDB: 6LNI). In this qualitative comparison, we notice that all three tools provide similar profiles and predict the amyloid fibril spanning residues 170−229. In Fig. 4B, we show the aggregation profile for human heterogeneous nuclear ribonucleoprotein D-like (UniProt: O14979) with the gray shaded area corresponding to the amyloid fibril spanning residues 345−395 (PDB: 7ZIR). Unlike Fig. 4A, in Fig. 4B, ZipperDB and AGGRESCAN do a poor job in predicting the amyloid fibril spanning residues 345−395, while the AmyloDeep prediction of that region is evident.

**Figure 4.**
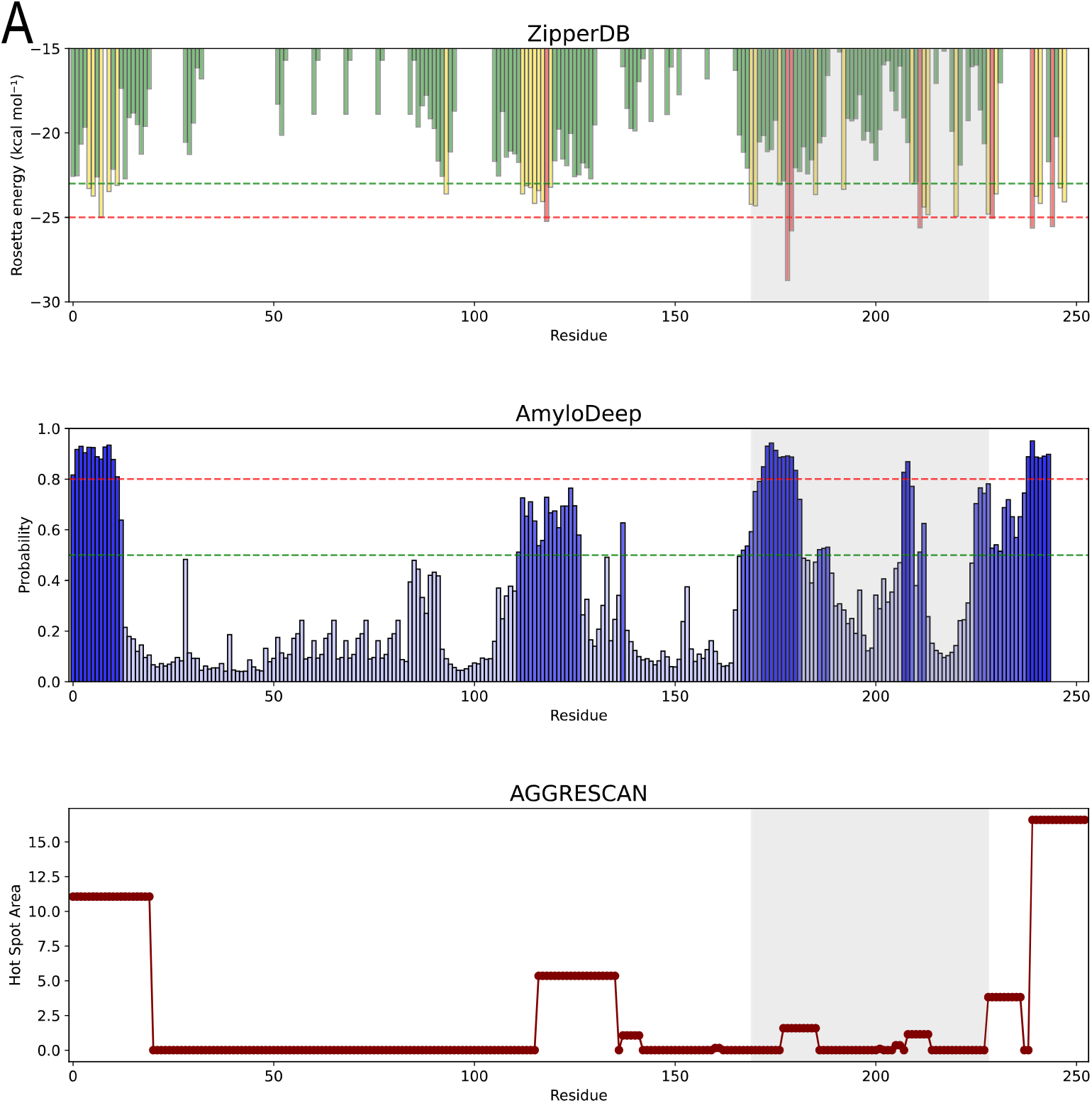
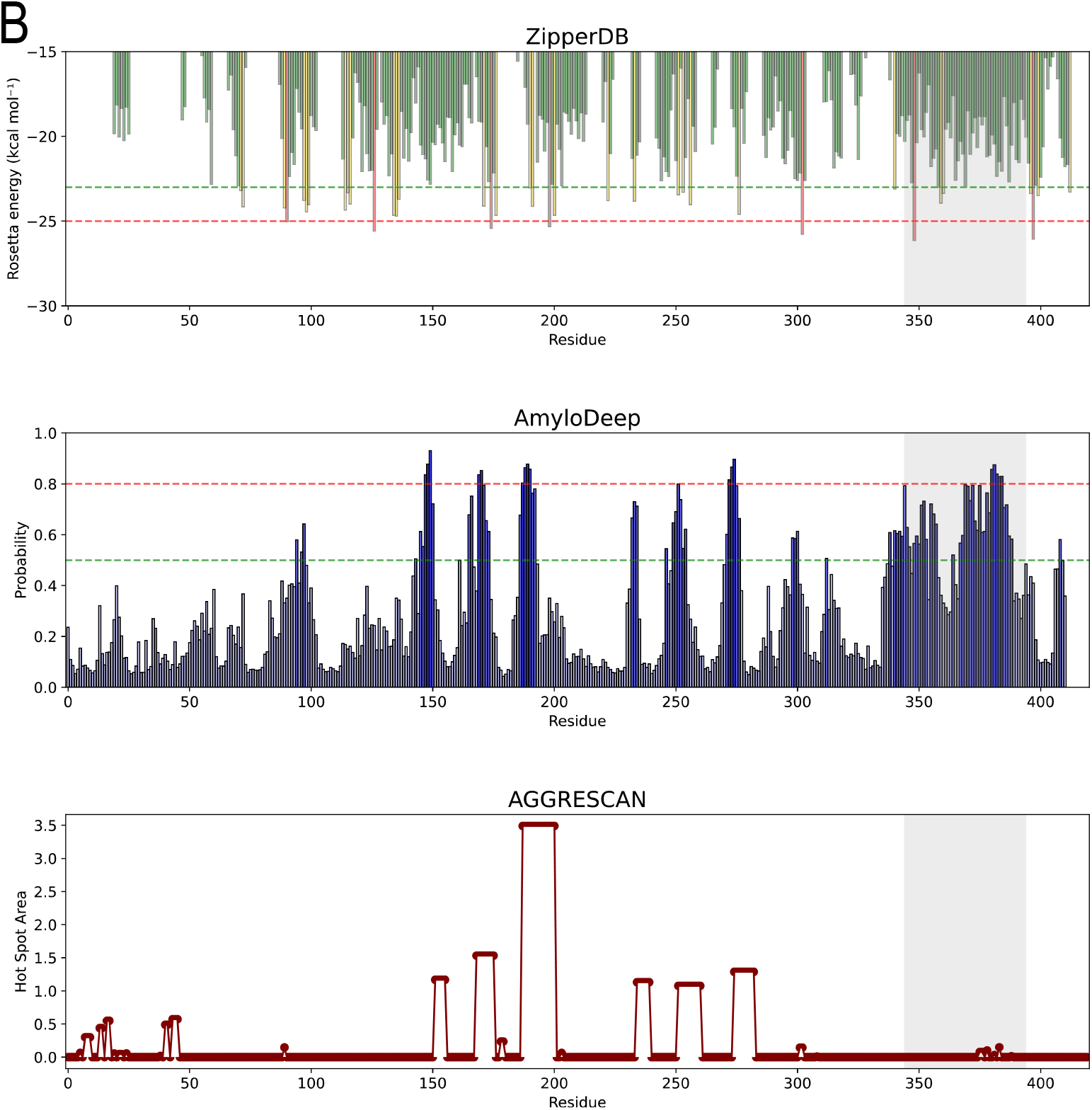
Qualitative comparison of AmyloDeep with well-established methods, ZipperDB (11), and AGGRESCAN (10) using aggregation profiles. A. We used the human PrP sequence (UniProt: P04156) for comparison. The gray shaded area corresponds to the amyloid fibril spanning residues 170−229 (PDB: 6LNI). B. We used human heterogeneous nuclear ribonucleoprotein D-like sequence (UniProt: O14979) for comparison. The gray shaded area corresponds to the amyloid fibril spanning residues 345−395 (PDB: 7ZIR). For ZipperDB, the green dashed line shows the threshold value of residues with a high beta-zipper formation propensity, the red dashed line indicates the high-aggregation propensity threshold, the yellow bars indicate residues with a higher than average aggregation propensity, and the red bars indicate aggregation-prone regions. For AmyloDeep with a 10 amino acids running window, the green and red dashed lines indicate the thresholds for the amino acids with higher than 0.5 and 0.8 probabilities of amyloid formation, respectively, and bars, correspondingly, are colored with darker shades of blue.

## Discussion

We first briefly describe how the methods compared in this study were developed. AGGRESCAN uses the aggregation propensity values per amino acid derived from experiments using single-point mutations at the central position of Aβ42 (10, 37). Then it uses a running window of size 5, 7, 9, or 11, depending on the input sequence length (10). The latest version AGGRESCAN4D (38) predicts pH-dependent aggregation and has the protein 3D structure as an input. ZipperDB is using Rosetta energy calculated based on hexapeptide zipper crystal structures, hence, ZipperDB has a running window size of 6 amino acids for aggregation profiles (11). The current updated version of ZipperDB is based on a neural network that generates scores for individual hexapeptide sequences (https://zipperdb.mbi.ucla.edu/info). Cross-Beta predictor is an Extra Trees classifier trained on the Cross-Beta DB database, a database based on cross-beta structures (26). It has a minimum running window size of 15 amino acids for the aggregation profile. AMYPred-FRL is a stacked ensemble model trained on the Amy dataset (24), where only sequences with 50 amino acids or more are present, and AMYPred-FRL currently does not provide an aggregation profile option. In contrast to, for example, ZipperDB or Cross-Beta predictor, we did not concentrate on specific amyloid features such as zipper crystal structure or cross-beta structure, but we used all the available to us labeled experimental data. We believe this broadens our tool’s capabilities.

Our experiments with AlphaFold 2 and 3 (6, 7) showed that they are doing poorly in predicting amyloid structures. There could be several reasons for this. First, although in recent years many amyloid fibril structures have been determined with CrioEM and other methods and deposited into PDB, we still have few such structures (we have around 81240 unique protein structures (https://www.rcsb.org/stats/growth/nr/matching-uniprot) and only 615 amyloid fibril structures (https://people.mbi.ucla.edu/sawaya/amyloidatlas/) as of August 1, 2025), hence little data for training. A single amino acid sequence may yield either a native-like globular structure or an amyloid fibril, hence, the same sequence can produce at least two drastically different structures. AmyloDeep and similar models could become a part of protein structure prediction tools, where the model could get amyloid formation probability from AmyloDeep, and use that information, for example, to provide two structures to the users for a given sequence: one globular native folded structure, and the other, an amyloid fibril structure.

We used three different embeddings (ESM2 150M, ESM2 650M, and UniRep) (33, 34) and five different predictive models. This gives an advantage in the sense that different encoders can grasp different features and properties of an amino acid sequence, which could be important for predicting amyloid formation. Using diverse model families (FNN, SVM, XGBoost, fine-tuned ESM2 150m) provides diversity in a similar sense as for the encoders mentioned above. Finally, the soft voting integrates their probabilities, producing a stable, consensus decision. The number and size of the models and encoders, on the other hand, are limited by available computation resources for training and inference. Inference time is a key user-facing metric. Among our five models, the fastest in training and inference is UniRep combined with FNN. This is because the embeddings of UniRep are smaller. Independent of the length of the amino acid sequence, it returns a 1900-dimensional vector, while the embeddings of ESM2 depend on the length of the amino acid sequence (it is one of the dimensions of the embedding matrix). Hence, we deploy UniRep combined with FNN as a light version on our web server.

The EMS3 model and its embeddings recently became publicly accessible, allowing for custom fine-tuning (31). We evaluated the feasibility of fine-tuning the open-access variant ESM3-sm-open-v1, which contains approximately 1.4 billion parameters. However, we found that ESM3-sm-open-v1 underperformed compared to ESM2 150M for our specific use case, so we concluded that ESM2 was the more effective choice for our case.

In summary, we developed a novel model for amyloid propensity prediction from the amino acid sequence that showed state-of-the-art performance and made it accessible through a web server and a Python package. Similar tools have been used for determining fibril-forming segments that nucleate aggregation, for further experimental studies (12, 39), and for the development of a database for proteome aggregation predictions (40). Further development could be the consideration of the external conditions, such as pH, on amyloid propensity. Such steps have already been taken (38). These steps can also be useful for protein structure prediction tools, as the structure of a protein also depends on external conditions.

## Acknowledgements

This work was supported by the HESC of Armenia (Research project № 23IRF-1F04). The authors would like to acknowledge networking support sponsored by the COST Actions CA21160 (ML4NGP) and CA21169 (DYNALIFE). We are thankful to the AUA 200 ChangeMakers.

