## Supplementary Information for "AmyloDeep: pLM-based ensemble model for predicting amyloid propensity from the amino acid sequence"

Below we provide the performance metrics and receiver operating characteristic (ROC) curves for the five individual models that, after probability calibration, became part of the ensemble model. For each model, 5-fold cross-validation was used, and the final performance metrics presented below are for amyloid class prediction averaged across all validation folds.

**Fine-tuned ESM2 150M**

| | Accuracy | Sensitivity | Specificity | F1 score | MCC |
|---|---|---|---|---|---|
| Test | 85.07 % | 87.44 % | 86.14 % | 86.78 % | 61.02 % |
| Validation | 82.54 % | 89.50 % | 79.02 % | 83.94 % | 65.57 % |
| Train | 85.01 % | 85.75 % | 85.35 % | 85.55 % | 69.97 % |

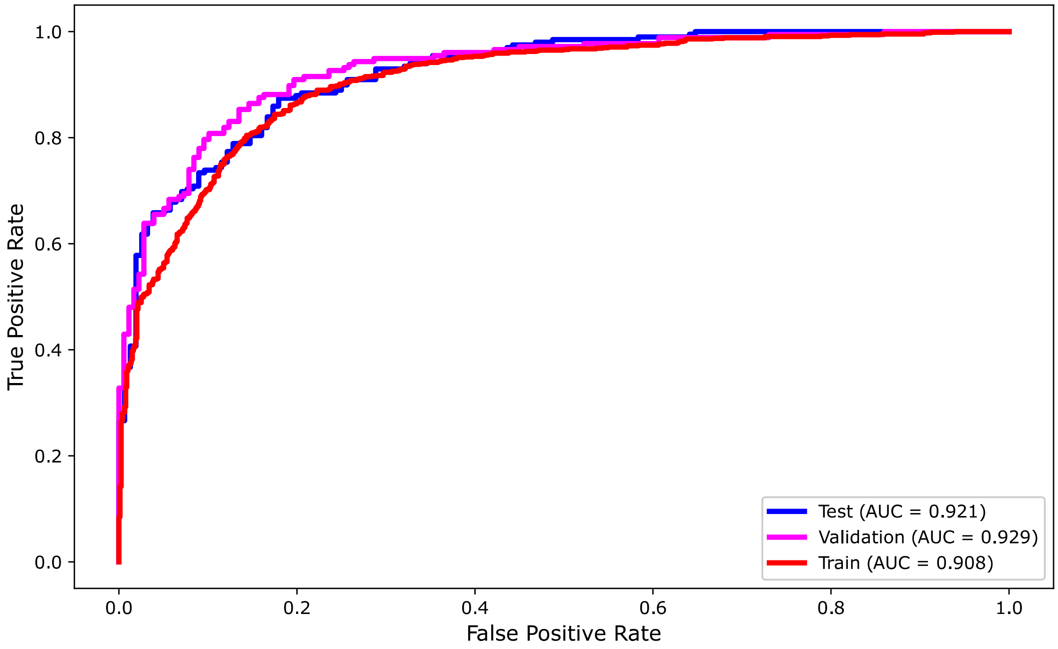


As mentioned in the methods section for ESM2-T30-150M, only the classification layer was fine-tuned, while the core parameters of the pretrained model remained unchanged. We have previously partially fine-tuned ESM2-T30-150M by unfreezing the last two encoder layers and the classification head. In that case model overfitted: training loss dropped quickly while validation performance declined, indicating memorization rather than generalization.

**UniRep combined with FNN**

| | Accuracy | Sensitivity | Specificity | F1 score | MCC |
|---|---|---|---|---|---|
| Test | 84.23 % | 85.43 % | 86.29 % | 86.86 % | 67.14 % |
| Validation | 84.79 % | 83.05 % | 85.96 % | 84.48 % | 70.32 % |
| Train | 87.55 % | 87.43 % | 88.46 % | 87.94 % | 75.12 % |

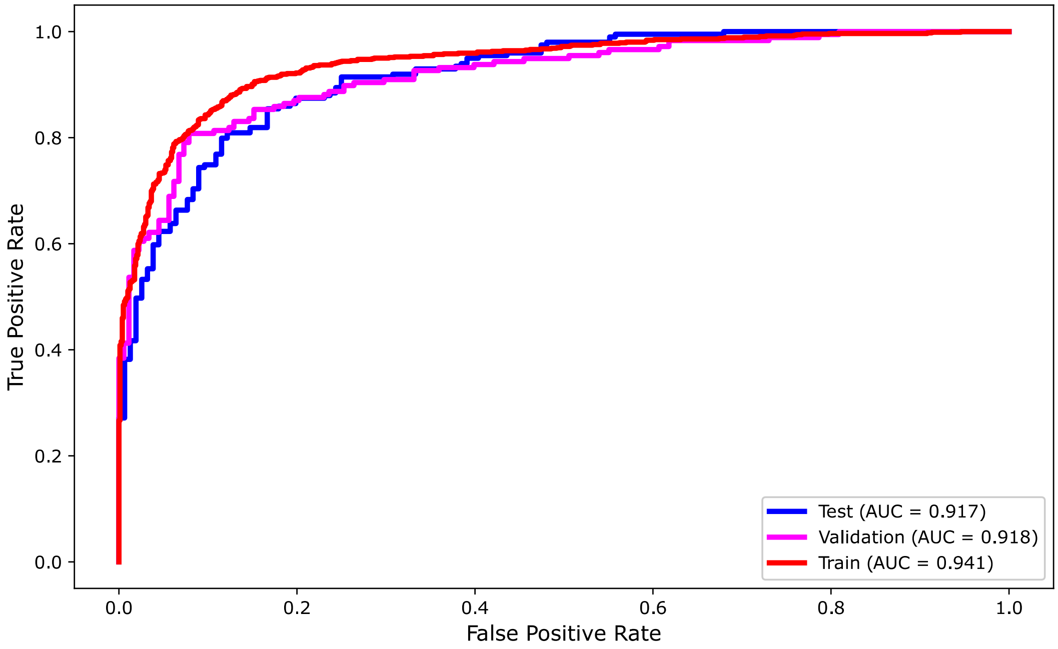

**ESM2 650M combined with FNN**

| | Accuracy | Sensitivity | Specificity | F1 score | MCC |
|---|---|---|---|---|---|
| Test | 81.41 % | 86.26 % | 84.78 % | 85.52 % | 62.45 % |
| Validation | 83.94 % | 81.91 % | 84.46 % | 84.03 % | 67.98 % |
| Train | 84.28 % | 81.91 % | 84.46 % | 83.16 % | 69.95 % |

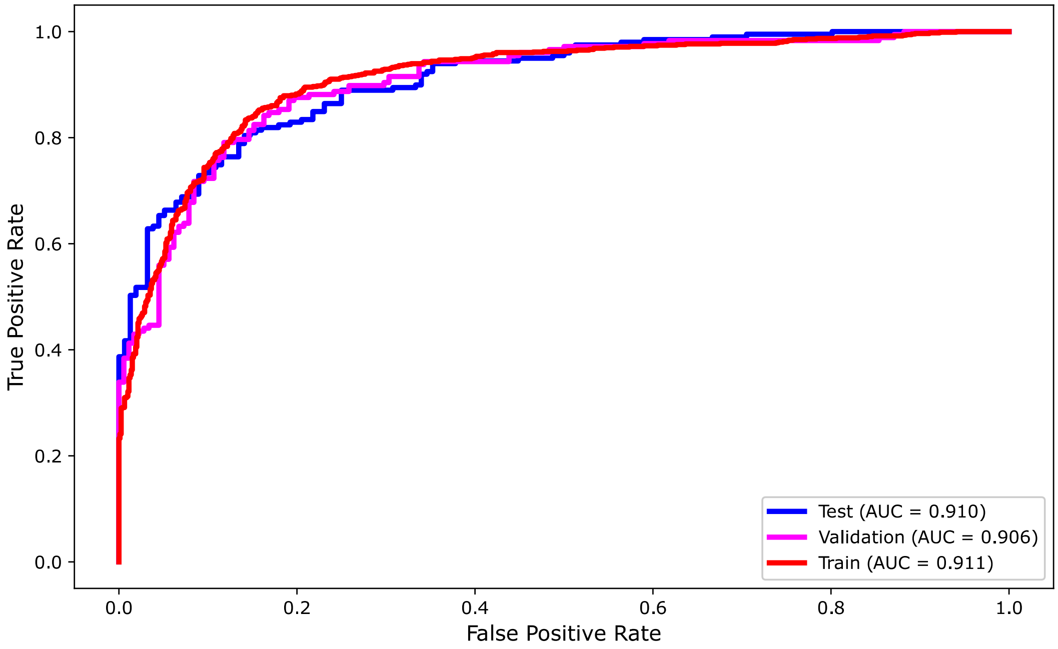

**ESM2 650M combined with SVM**

| | Accuracy | Sensitivity | Specificity | F1 score | MCC |
|---|---|---|---|---|---|
| Test | 84.23 % | 84.92 % | 86.67 % | 85.79 % | 68.08 % |
| Validation | 84.51 % | 84.18 % | 84.66 % | 84.42 % | 69.01 % |
| Train | 85.91 % | 85.89 % | 84.52 % | 86.54 % | 71.77 % |

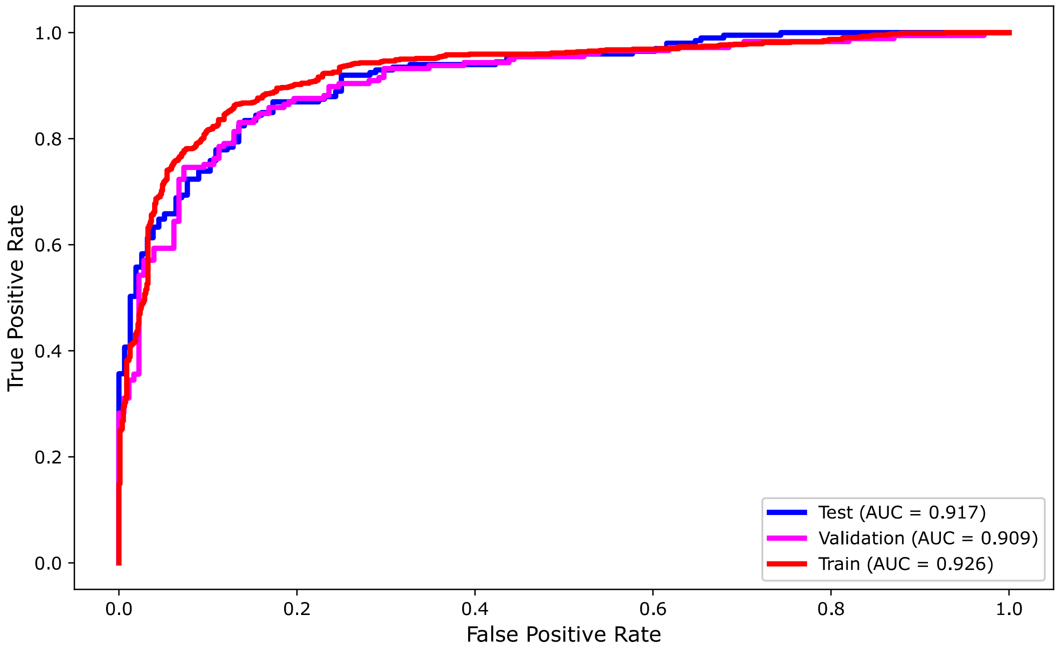

**ESM2 650M combined with XGBoost**

| | Accuracy | Sensitivity | Specificity | F1 score | MCC |
|---|---|---|---|---|---|
| Test | 83.38 % | 86.43 % | 84.31 % | 85.36 % | 66.17 % |
| Validation | 83.10 % | 88.14 % | 80.00 % | 83.87 % | 66.54 % |
| Train | 84.89 % | 88.82 % | 83.21 % | 85.92 % | 69.81 % |

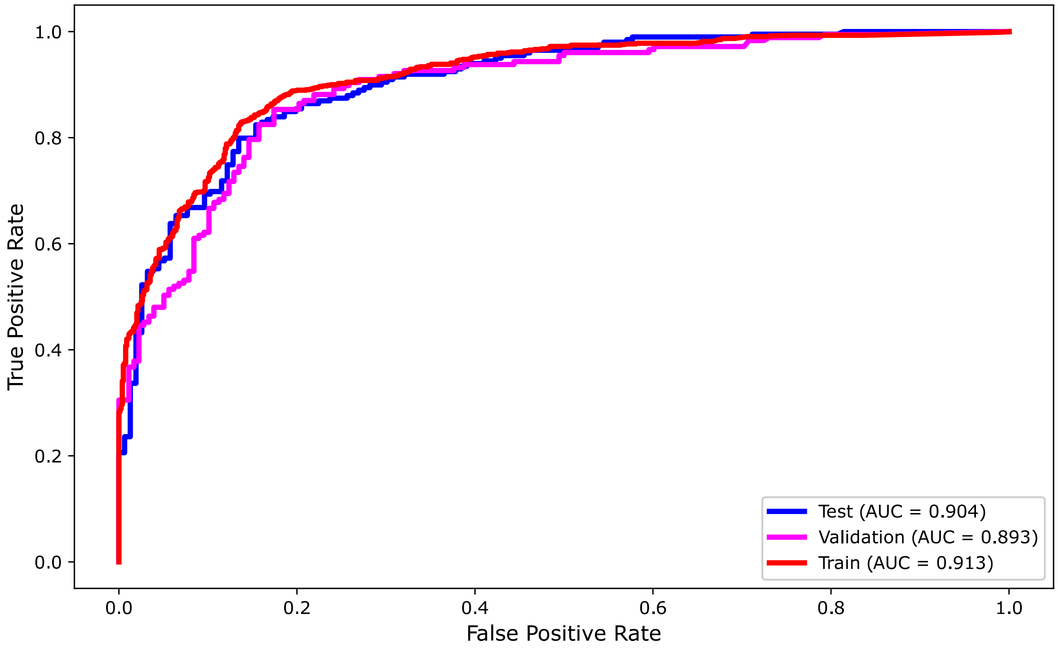
